# Haploinsufficiency of the psychiatric risk gene *Cyfip1* causes abnormal postnatal hippocampal neurogenesis through microglial and Arp2/3 mediated actin dependent mechanisms

**DOI:** 10.1101/417832

**Authors:** Niels Haan, Laura J Westacott, Jenny Carter, Michael J Owen, William P Gray, Jeremy Hall, Lawrence S Wilkinson

**Affiliations:** Neuroscience and Mental Health Research Institute and MRC Centre for Neuropsychiatric Genetics and Genomics, Cardiff University, Hadyn Ellis Building, Maindy Road, Cardiff, United Kingdom; Hodge Centre for Neuropsychiatric Immunology, Cardiff University, Hadyn Ellis Building, Maindy Road, Cardiff, United Kingdom; Brain Repair and Intercranial Neurotherapeutics Unit, Cardiff University, Hadyn Ellis Building, Maindy Road, Cardiff, United Kingdom; School of Psychology, Cardiff University, Tower Building, Cardiff, United Kingdom

**Author notes:** Corresponding authors: Dr Niels Haan.

**Keywords:** Cyfip1, microglia, apoptosis, neurogenesis, schizophrenia

## Abstract

Genetic risk factors can significantly increase chances of developing psychiatric disorders, but the underlying biological processes through which this risk is effected remain largely unknown. Here we show that haploinsufficiency of Cyfip1, a candidate risk gene present in the pathogenic 15q11.2(BP1-BP2) deletion may impact on psychopathology via abnormalities in cell survival and migration of newborn neurons during postnatal hippocampal neurogenesis. We demonstrate that haploinsufficiency of *Cyfip1* leads to increased numbers of adult born hippocampal neurons due to reduced apoptosis, without altering proliferation. We confirm this is due to a cell autonomous failure of microglia to induce apoptosis through the secretion of the appropriate factors. Furthermore, we show an abnormal migration of adult-born neurons due to altered Arp2/3 mediated actin dynamics. Together, our findings throw new light on how the genetic risk candidate *Cyfip1* may influence the hippocampus, a brain region with strong evidence for involvement in psychopathology.

## Introduction

Many psychiatric conditions show high heritability. Recent genetic studies in schizophrenia for example have identified up to 160 loci that increase risk for the disease (Rees *et al*., 2014; Pardiñas *et al*., 2018) enriched within synaptic (Hall *et al*., 2015), histone modifying and immune system genes(Network and Pathway Analysis Subgroup of Psychiatric Genomics Consortium, 2015). *CYFIP1* is a genetic risk factor for schizophrenia, autism and developmental delay by virtue of its presence in the highly penetrant 15q11.2(BP1-BP2) copy number deletion (Cox and Butler, 2015). Loss of one copy of this interval leads to substantially increased risk for disorder in both Western (Stefansson *et al*., 2008; Kirov *et al*., 2009) and Han Chinese populations (Zhao *et al*., 2013). *CYFIP1* haploinsufficiency is likely a major contributor to the 15q11.2(BP1-BP2) psychiatric phenotype due to evidence of *CYFIP1*’s involvement in a range of synaptic functions, including key roles in dendritic spine morphology and branching (Pathania *et al*., 2014; Oguro-Ando *et al*., 2015; Hsiao *et al*., 2016). CYFIP1 protein influences synaptic function in two main ways. CYFIP1 interacts with fragile X mental retardation 1 (FMRP), the protein gene product of FMR1to suppress translation of up to 800 target transcripts (Darnell *et al*., 2011) and also with Wiskott-Aldrich syndrome protein family member 1 (WAVE1), a key mediator of cytoskeleton dynamics, to modulate ARP2/3 dependant actin branching (Chen *et al*., 2010)

In the present work we focus on microglia and adult hippocampal neurogenesis (AHN) as important sites of CYFIP1 action and as potential contributors to psychiatric phenotypes arising from 15q11.2(BP1-BP2) copy number deletion. Adult born hippocampal neurons, up to 700 of which are estimated to be born per day in humans (Spalding *et al*., 2013), develop through a well-defined series of cellular events, starting from proliferative stem cells and intermediate progenitors, through migratory neuroblasts and immature neurons, before eventually forming mature granule cells which distribute through the dentate gyrus (Kempermann, Song and Gage, 2015). In rodent models, adult born hippocampal neurons have been shown to be involved in a number of behavioural processes spanning pattern separation (Clelland *et al*., 2009; Sahay *et al*., 2011), spatial navigation (Kee *et al*., 2007; Dupret *et al*., 2008), memory turnover (Akers *et al*., 2014), and acquisition (Seo *et al*., 2015) and extinction (Deng *et al*., 2009) of fear memory. These behaviours are affected in psychiatric disorder including schizophrenia (Holt *et al*., 2009; Das *et al*., 2014; Martinelli and Shergill, 2015; Salgado-Pineda *et al*., 2016). Antipsychotic drugs, specifically atypical antipsychotics, also impact AHN (Newton and Duman, 2007) and importantly, a number of functionally diverse schizophrenia risk genes, including *DISC1* (Ye *et al*., 2017), *SREB2/GPR85* (Chen *et al*., 2012), *CACNA1C* (Temme *et al*., 2016; Moon *et al*., 2018) *DGCR8* (Ouchi *et al*., 2013), and *miR137* (Smrt *et al*., 2010), converge in their ability to modify AHN. Additionally, post-mortem findings from schizophrenia patients reveal changes in cell proliferation and expression of immature neuronal markers in the hippocampus (Barbeau *et al*., 1995; Reif *et al*., 2006; Allen, Fung and Shannon Weickert, 2016).

Immune changes are another common hallmark of several psychiatric disorders, including schizophrenia where evidence from epidemiological data (Brown and Derkits, 2010; Knuesel *et al*., 2014) and findings showing increased inflammatory factors in the circulation (Miller *et al*., 2011), and in post-mortem brain (Trépanier *et al*., 2016) of schizophrenia patients, implicate altered immune function in the development of disease symptoms. Mechanistic insights linking immune changes to risk for psychopathology are however largely lacking. Recent interest has focused on microglia, the resident macrophage-like immune cells of the central nervous system, with data suggesting possible alterations in microglial functioning in both those at high risk and those diagnosed with schizophrenia (Bloomfield *et al*., 2016; Holmes *et al*., 2016; Notter, Coughlin, Gschwind, *et al*., 2018; Notter, Coughlin, Sawa, *et al*., 2018). The causal mechanism(s) by which microglia might contribute to pathogenesis remain unclear, though there is increasing evidence that, in addition to canonical immune functions, microglia also modify neuronal networks through synaptic pruning during development (Paolicelli *et al*., 2011) and into adulthood (Schafer *et al*., 2012). This microglial-mediated pruning is important for normal brain and behavioural functioning (Kim *et al*., 2017). Additionally, microglia can modulate neuronal firing through direct cell-cell contacts (Li *et al*., 2012), and have been shown to be capable of phagocytosing both developing (Cunningham, Martinez-Cerdeno and Noctor, 2013) and adult (Brown and Neher, 2014) neurons.

In this work we used a translationally relevant haploinsufficient mouse model to reveal hitherto undescribed effects of manipulating *Cyfip1* on the survival and migration of newborn neurons in hippocampus, key cellular processes in the development and function of the adult hippocampus and associated circuitry. We also reveal the mechanisms mediating these effects, highlighting CYFIP1 effects on microglia mediated apoptosis and Arp2/3 dependent actin dynamics, respectively. We suggest these findings may have relevance to how *CYFIP1* haploinsufficiency can influence hippocampal function in the context of the markedly increased risk for psychiatric disorder arising from the 15q11.2(BP1-BP2) copy number deletion.

## Materials and methods

For full materials and methods, see Supplementary Information.

### *Cyfip1* heterozygous knockout mice

Cyfip1^tm2a(EUCOMM)Wtsi^ animals (MGI:5002986, referred to hereafter as *Cyfip1*^+/-^) were maintained heterozygously on a C57/BL6J background. Animals used were of mixed sex and 8-12 weeks old. *Cyfip1*^+/-^ and wildtype littermates were kept in conventional cages with 12h light-dark cycle and ad libitum access to water and food at all time.

### Tissue harvesting and immunohistochemistry

Paraformaldehyde fixed brains from 8-12 week old *Cyfip1*^+/-^ and wildtype littermates were prepared as previously described (Haan *et al*., 2013). Free floating 40 μm cryostat sections were used for immunohistochemistry. Immunohistochemistry was performed as previously described(Haan *et al*., 2013), with a 1:12 stereotactic sampling rate. Detection of cleaved caspase 3 used a modified protocol, where primary antibodies were incubated for 48 h in blocking solution at 4°C. Primary and secondary antibodies are described in table S1.

### Primary hippocampal progenitor culture

Hippocampal progenitors were isolated from P7-P8 animals. Isolated hippocampi were dissociated in papain and progenitors were enriched through Optiprep density gradient centrifugation. Cells were cultured on poly-L-lysine and laminin in the presence of EGF and FGF2.

### Primary microglial culture

Microglia were isolated from P7-P8 whole brain mixed glia through the shake-off method.

### Immunocytochemistry

Fixed cultures were blocked and permeabilized with 5% donkey serum and 0.1% Triton X100 in PBS for 30 minutes at room temperature. Primary antibodies were applied in the same solution overnight at 4 °C. Antibodies are detailed in Table S1. Following three washes in PBST, relevant secondary antibodies were applied in PBST for two hours at room temperature. For identification of microglia, Alexa 568 conjugated IB4 (Thermo Fisher I21412) was added to secondary antibody solutions at 2.5 μg/ml. Cells were washed, counterstained with DAPI, and mounted for microscopy.

### Conditioned medium

Microglia conditioned medium was prepared by incubating 5×10^5^ microglia per ml in progenitor medium for 24 h. 25% conditioned medium was added to hippocampal cultures.

### Time-lapse imaging and cell tracking

Primary hippocampal progenitors were plated on poly-D-lysine and laminin coated 24 well plates, and cultured for 5 days. At 5 DIV, cultures were imaged on an inverted Leica DMI600B microscope, at 37 °C, with 5% CO_2_ in air. Cells were imaged for two hours, with image acquisition every 5 minutes. Total distance covered was determined through manual cell tracking using the MTrackJ plugin in ImageJ (Meijering, Dzyubachyk and Smal, 2012). A minimum of 50 cells ware analysed per condition per animal.

### Actin analysis

Cells were fixed at 5 DIV with 4% paraformaldehyde in PBS, at 4 °C for 30 minutes, permeabilised with 0.1% Triton X100 in PBS for 15 minutes, and incubated with 300 nM Alexafluor 488 conjugated DNAseI (ThermoFisher D12371) and 200 nM Alexafluor 647 conjugated phalloidin (ThermoFisher A22287) for 30 minutes. Fluorescence intensity was measured in ImageJ.

### Arp2/3 inhibition

Cells were treated with 50 μM CK-548 or 250 μM CK-666 in DMSO, or vehicle, for two hours prior to time-lapse imaging. Drugs were present during imaging.

## Results

### *Cyfip1*^+/-^ animals have normal proliferation but increased numbers of adult born immature neurons in the hippocampus

We first investigated cell proliferation in the dentate gyrus of the hippocampus. Animals received a single injection (i.p) of 100 mg/kg BrdU 6 hours prior to sacrifice. No significant differences in cells positive for BrdU (197.3±11.1 vs 182.9±11.7 cells/mm^2^, t_(7)_=0.86, p=0.41) or the cell cycle marker Ki67 (422.4±24.9 vs 371.8±30.8 cells/mm^2^, t_(7)_=1.27, p=0.25) were found (fig 1A-C), indicating no effects on cell proliferation. However, staining for the early neuronal marker doublecortin (DCX) revealed a significant increase of immunoreactive cells in the *Cyfip1*^+/-^ animals (367.0±21.9 vs 550.6±39.4 cells/mm^2^, t_(15)_=4.38, p=6.21×10^-4^, fig 1D-E), consistent with the presence of increased numbers of immature neurons. In a 30 day BrdU pulse-chase protocol we observed increased numbers of mature post-mitotic BrdU+/NeuN+ labelled cells in the dentate gyrus of *Cyfip1*^+/-^ animals (42.9±7.6 vs 61.5±3.9 cells/mm^2^, t_(10)_=4.54. p=6.8×10^-4^), as well as an increased fraction of NeuN+ cells in the total BrdU+ population (51.0±2.7% vs 60.5±0.8%, t_(10)_=3.95, p=0.002) (fig 1F-H). These data indicated that the excess numbers of immature neurons seen in the *Cyfip1*^+/-^ animals were maintained into maturation. We found no significant effects of sex of the animal (see Supplementary Results and fig S2).

**Figure 1:**
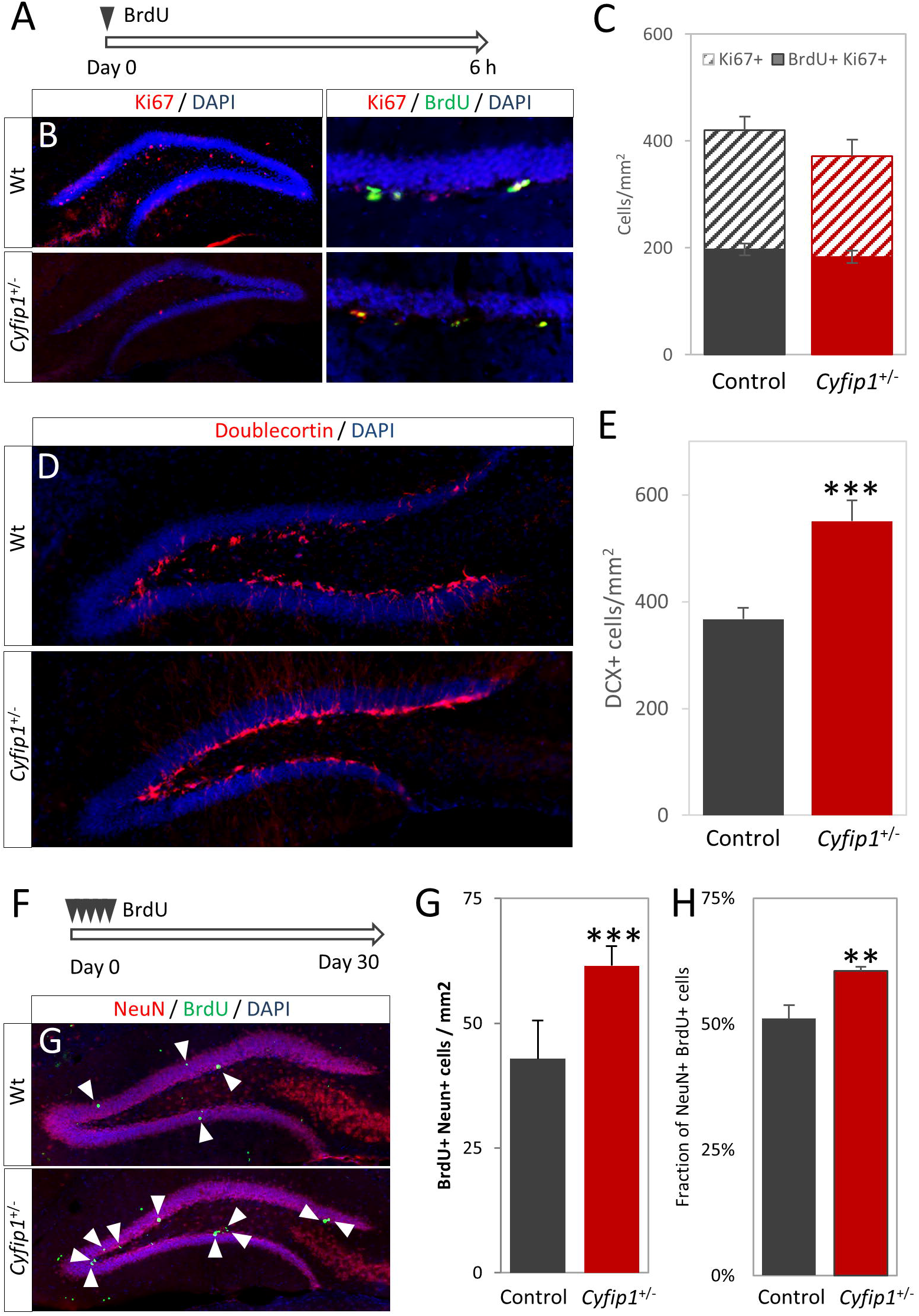
Cyfip1 haploinsufficiency increases neurogenesis in vivo in the absence of changes in proliferation. a) *Experimental scheme*, proliferation rates were studied with a 6h pulse of BrdU. b, c) Representative immunohistochemistry showing distribution and location of proliferative (Ki67+) and dividing (Ki67+BrdU+) cells in wild-type and *Cyfip1*^+/-^ animals (n=4 each), respectively. d) Quantification showed no difference in numbers of proliferative cells or number of cell divisions. e, f) Representative staining for DCX in wild-type and *Cyfip1*^+/-^ animals (n=8 each), respectively. g) Quantification showed a significantly larger number of immature neurons in the *Cyfip1*^+/-^ animals. h) *Experimental scheme*, cell maturation was studied with a 30 day BrdU pulse-chase. i,j) Representative images showing BrdU÷ and NeuN+ cells. Arrowheads indicate BrdU+ cells k) *Cyfp1*^+/-^ animals (n=6) had significantly higher numbers of BrdU+NeuN+ cells than the wild-type animals (n=5). l) The fraction of BrdU+ cells which had matured into NeuN+ cells was significantly increased in the *Cyfip1*^+/-^ animals. All data are shown as mean±SEM.

### Increased neuronal numbers in *Cyfip1* haploinsufficiency is due to decreased apoptosis

To investigate the cell-intrinsic effects of *Cyfip1* haploinsufficiency independent of niche effects we used primary hippocampal progenitor cultures. The *in vitro* data corroborated the ex *vivo* finding, showing an increase in the fraction of DCX immunoreactive cells in the cultures prepared from *Cyfip1*^+/-^ animals (5.5±0.5% vs 8.8±1.4%, t_(16)_=2.78, p=0.014, fig 2A,B) in the absence of any effects on proliferation (see Supplementary Results and fig S3). There were no differences in the proportions of earlier progenitors of the 1/2a GFAP+/nestin+ type (9.4±1.2% vs 10.0±1.7%, t_(35)_=0.76, p=0.79), or 2b GFAP-/nestin+/DCX-types (22.1±1.6% vs 20.2±2.5%, t_(35)_=0.67, p=0.51), (fig 2C-E), indicating no effects on early stages of neurogenesis. We next assessed general cell viability and survival. We saw a significant increase in the proportion of viable MitoTracker+ cells (82.4±1.2% vs 89.6±0.7%, t_(14)_=3.23, p=0.002) and fewer propidium iodine positive dead/dying cells (7.7±0.6% vs 5.4±0.5%, t_(16)_=2.44, p=0.026)(fig 2F-H). We then investigated apoptosis specifically in the DCX+ populations. We found a marked reduction in the fraction of DCX+ /nestin-cells positive for cleaved-caspase-3 (25.9±5.5% vs 7.2±2.5%, t_(16)_=2.32, p=0.034, fig 2I,J). We found no sex effects in any of the phenotypes investigated in the primary cultures (see Supplementary Results and fig S4). Together, these data suggested *Cyfip1* haploinsufficiency impaired a normal homeostatic mechanism for modulating newborn neuron number, leading to a greater number surviving to go on to maturity. To confirm the effects on cell death due to apoptosis were not an artefact of the culture system we also showed that there were significantly fewer cleaved-caspase-3 immunoreactive cells *in vivo* in the dentate gyrus of *Cyfip1*^+/-^ animals (2.1±0.2 vs 1.2±0.1 cells/mm^2^, t_(23)_=3.01, p=0.006, fig 2K,L), demonstrating a high level of concordance between effects in cell culture and intact brain tissue.

**Figure 2:**
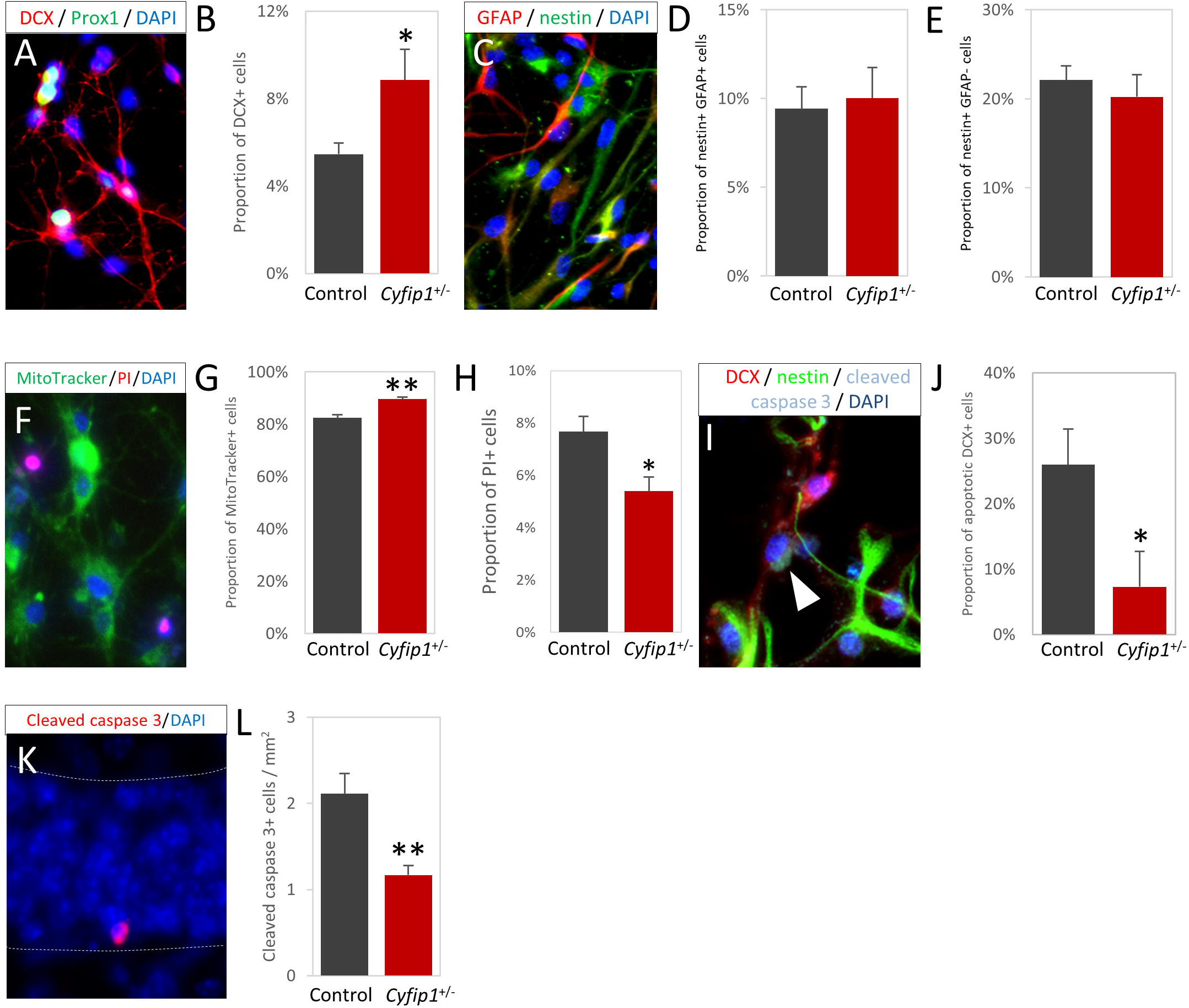
Primary hippocampal progenitor cultures prepared from P7-P8 brain confirm ex vivo findings and indicate that increased neuronal numbers in Cyfip1^+/-^ animals is due to reduced apoptosis. a) Immunoreactivity for immature neuron marker DCX and hippocampal marker Prox1 confirm cultures contain hippocampus-derived developing neurons. b) Cyfip1^+/-^ animals showed a significantly higher proportion of DCX+ cells in the total DAPI+ population (n=17). c) Representative staining for progenitor markers GFAP and nestin showed cultures contained a mixed population of progenitors at differing stages of development. d) There was no significant difference in the proportion of early GFAP+/nestin+ cells (n=36). e) The proportion of later GFAP-/nestin+ cells (n=36) was also unaffected. f. Live imaging showed cultures are comprised of healthy, MitoTracker+ cells, with some dying PI+ cells. g) A small but significant increase in the proportion of MitoTracker+ cells was observed in the *Cyfip1*^+/-^ cultures. (n=15) h) Conversely, a significant decrease in the proportion of PI+ cells was seen in the *Cyfip1*^+/-^ cultures (n=14). i) Immunoreactivity for apoptotic marker cleaved caspase with DCX and nestin showed specific neuronal apoptosis. Arrowhead indicates an apoptotic neuron. j) The fraction of neurons undergoing apoptosis was significantly lower in the *Cyfip1*^+/-^ cultures (n=17). I) Representative image of apoptotic cells in the detate gyrus. L) As seen *in vitro, Cyfip1*^+/-^ animals (n=13) have significantly fewer apoptotic cells than wild type animals (n=12). All data are shown as mean±SEM.

### Evidence that apoptosis of newly born neurons is mediated by microglia secreted factors

Given the known roles microglia play in earlier stages of neurogenesis (Ziv *et al*., 2006), we speculated that microglia may also play a role in the survival and death of adult new born immature neurons, and further that they may play a role in the effects of *Cyfip1* haploinsufficiency. In a series of preliminary assessments we first established there were no genotype differences in the density of Iba1+ microglia in hippocampus ex vivo (177.1±4.1 vs 162.9±8.7, t_(12)_=1.22, p=0.25, fig 3A,B), or in vitro (fig 3C,D) or effects of sex on this measure (supplementary data and fig S5. We next showed that Cyfip1 is highly expressed in primary wildtype microglia (comparable to the levels of the housekeeping gene GAPDH) and in wild type cells expression is strongly up-regulated (15.2±0.95-fold) following activation with lipopolysaccharide (LPS), (t_(5)_=14.8, p=0.001, fig 3E), suggesting Cyfip1 is involved in microglial regulation. Then we compared LPS-activation of wildtype and *Cyfip1*^+/-^ primary microglia indexed by CD68 immunoreactivity. Both cultures showed evidence of a robust activation with a significant treatment effect (F_(1,48)_=9.86, p=0.002). There was also a significant genotype effect (F_(1,48)_=4.27, p=0.043) with no interaction (F_(1,48)_=0.93, p=0.33) reflecting a degree of blunting of activation in *Cyfip1*^+/-^ cells, (fig 3F).

**Figure 3:**
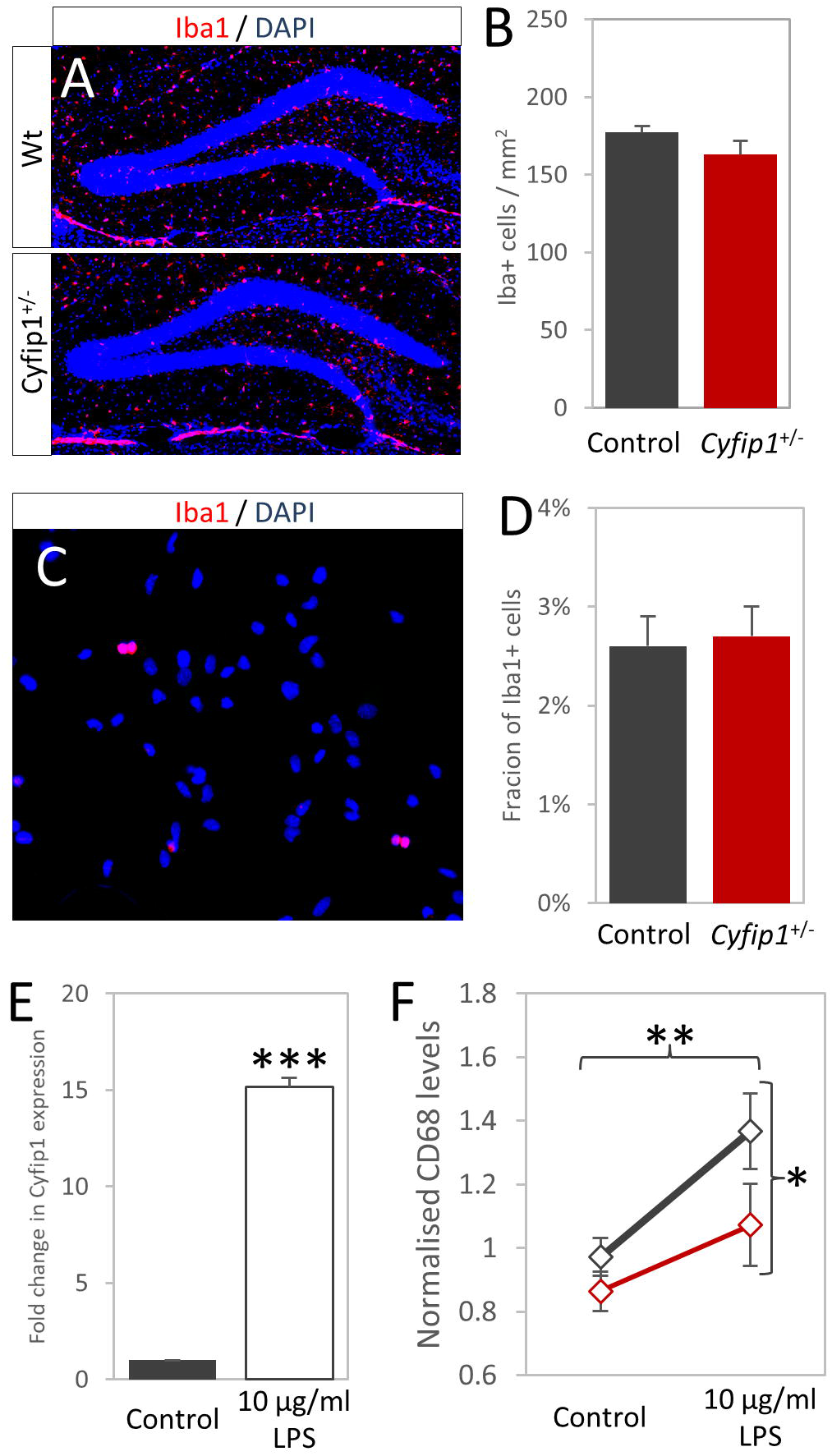
Microglia function, but not number is affected by Cyfip haploinsufficiency. a) Immunocytochemisty for Iba1 shows the numbers of microglia in the dentate gyrus. b) Quantification shows no effects of genotype on the density or microglia. c) Iba1 immunocytochemistry in primary hippocampal cultures shows low propositions of microglia present. d) There are no significant difference between genotypes in the Iba1 proportions.

To investigate the role of microglia in neuronal apoptosis, we depleted microglia from wildtype primary hippocampal progenitor cultures using the specific microglia toxin Mac-1-SAP, targeting CDllb expressing cells having first established the efficacy of the toxin against primary mouse microglia (fig 4A). Depletion of microglia from hippocampal cultures led to an increase in the proportion of DCX+ cells (7.5±1.6% vs 12.7±0.3%, t()=3.19, p=0.019, fig 4B), and a specific decrease in the fraction of apoptotic DCX+ cells (13.1±3.1% vs 2.7±1.7%, t(7)=2.93, p=0.026, fig 4C), without any associated changes in the nestin+ population (50.7±2.7 %vs 53.8±1.0%, t(7)=1.07, p=0.32, fig 4D), phenocopying the *Cyfip1*^+/-^ cultures, and suggesting a role for microglia in regulating DCX numbers. Conversely, addition of exogenous microglia on a semi-permeable membrane, allowing only for soluble factors to influence neuronal apoptosis, had opposite effects. Under these conditions we observed a decreased proportion of DCX+ cells (5.9±0.8% vs 1.9±0.5%, t_(15)_=4.36, p=6.5×10^-4^, fig 4F), increased apoptosis in the DCX+ population (14.8±4.7% vs 51.7±9.0%, t_(15)_=3.27, p=0.006, fig 4G) and no effect on the nestin+ population (58.4±3.3% vs 62.7±3.0%, t_(15)_=0.98, p=0.34, fig 4H). These experiments provided converging evidence for a previously unknown function of microglia in inducing apoptosis of immature neurons in hippocampus mediated, at least in part, by soluble factors.

**Figure 4:**
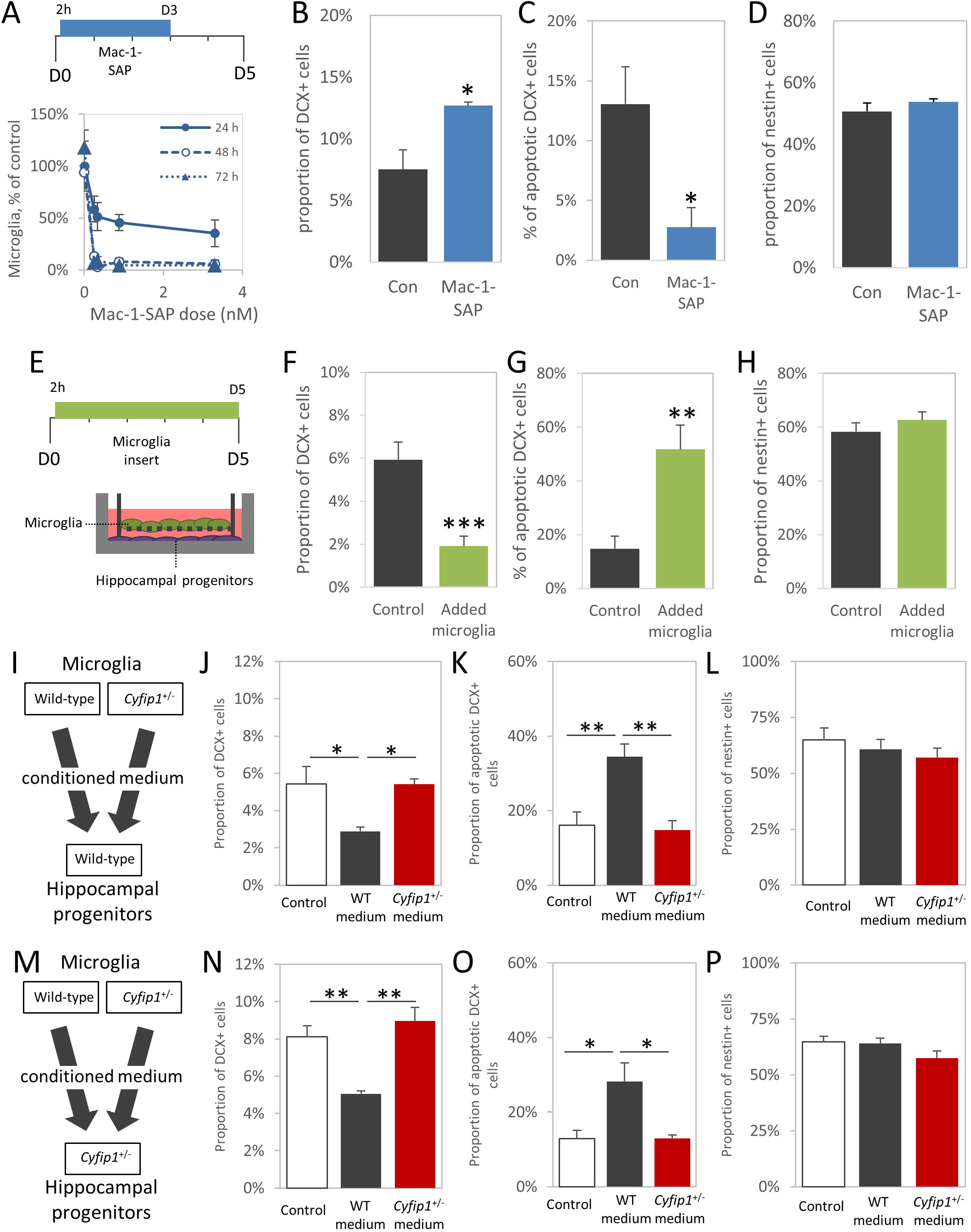
Cyfip1 *haploinsufficiency affects the ability if microglia to regulate number and apoptosis rates of immature neurons*. a) *Experimental setup*, Mac-1-Sap was present in wildtype hippocampal progenitor cultures from 2 hours after isolation until 3 days *in vitro*. Dose-response curves of microglial death after Mac-1-SAP administration show near-complete depletion of Iba1+ microglia was achieved within 48 h (n=6) b). Cultures where microglia were depleted (n=4) showed a higher proportion of DCX+ cells than controls (n=4). c) Correspondingly, the fraction of DCX cells undergoing apoptosis, as marked by cleaved caspase 3 staining, was significantly decreased in the absence of microglia, d) The proportion of nestin+ cells was unaffected by microglia depletion. e) *Experimental setup*, Membrane inserts containing wild-type microglia prepared from P7-8 brain were added to cultures 2 hours after progenitor isolation and were present throughout the remainder of the experiment. Due to the experimental conditions, no cell-cell contact between microglia and progenitors was possible, but secreted factors were free to diffuse. f) The presence of microgliacontaining inserts (n=8) significantly decreased the proportion of DCX expressing cells compared to controls (n=8). g) Conversely, the added microglia significantly increased apoptosis in DCX+ cells. H) No effect was seen in nestin expressing cells. i) *Experimental design*, Wild-type hippocampal progenitors were exposed to conditioned medium from either wildtype or *Cyfip1*^+/-^ microglia, j) In wildtype cultures, conditioned medium from wildtype microglia induced a significant reduction in the proportion of DCX+ cells, whereas medium from *Cyfip1*^+/-^ microglia was unable to do this (n=4 in all conditions). k) An inverse pattern was seen in neuronal apoptosis where wildtype microglia medium increased apoptosis in DCX+ immature neurons but *Cyfip1*^+/-^ medium did not. I) Nestin+ cells were not significantly affected by the presence of microglial factors. m) *Experimental design, Cyfip1*^+/-^ hippocampal progenitors were exposed to conditioned medium from either wildtype or *Cyfip1*^+/-^ microglia. n) As seen in the wildtype cultures, a significant decrease in DCX+ cells was observed only after stimulation with medium from wildtype microglia (n=4 in all conditions). o) Similarly, medium from *Cyfip1*^+/-^ microglia was unable to induce apoptosis in *Cyfip1*^+/-^ progenitors. p) Again, the nestin* population was unaffected. All data are shown as mean±SEM.

### Haploinsufficiency of *Cyfip1* inhibits the ability of microglia to support apoptosis of new born immature hippocampal neurons

Having established a novel microglial dependent mechanism for regulation of neuronal survival we next tested whether effects on this process could underlie the effects of *Cyfip1* haploinsufficiency on neuronal survival. To directly address the potential role the induction of apoptosis by microglial secreted factors in an unbiased way, we used a cross-genotype conditioned medium approach. Medium obtained from wildtype and *Cyfip1*^+/-^ microglia preparations was added to both the wildtype and *Cyfip1*^+/-^ hippocampal progenitor enriched cultures (see fig 4I and M. for schematic of design) and measured cell proportions and levels of apoptosis. In the wildtype progenitor cultures adding conditioned medium from wildtype microglia resulted in a significant reduction in the proportion of DCX+ cells, compared to unstimulated cells, whereas *Cyfip1*^+/-^ conditioned medium had no effect (5.4±0.9% vs 2.9±0.2% vs 5.4±0.3% for unstimulated, wildtype stimulated and *Cyfip1*^+/-^ stimulated respectively, main effects of genotype F_(2,9)_=6.31, p=0.019, fig 4J). *Post hoc* testing confirmed significant differences between controls and wildtype medium (p=0.031) and between those stimulated with wildtype and *Cyfip1*^+/-^ medium (p=0.033). This pattern of effects was mirrored inversely in the fraction of DCX+ cells that were also immunopositive for cleaved-caspase-3, with only conditioned medium from wildtype microglia cultures increasing apoptosis (16.1±3.6% vs 34.5±3.5% vs 14.8±2.5%, main effects of genotype F_(2,9)_=11.57, p=0.003, fig 4K). Again, *post hoc* testing showed the wildtype stimulated condition was significantly different from controls and *Cyfip1*^+/-^ stimulated condition (p=0.007 and p=0.005, respectively). Consistent with the conclusion that these effects were limited to the DCX+ immature neuron population of cells, the proportion of cultures that was nestin+ was unchanged across all conditions (65.0±5.2% vs 60.7±4.4% vs 57.1±4.1%, F_(2,9)_=0.75, p=0.50, fig 4L).

Importantly, identical patterns of effects were seen in the *Cyfip1*^+/-^ hippocampal progenitor cultures. Using an identical experimental design (fig 4I), we found that conditioned medium from wildtype microglia, but not from *Cyfip1*^+/-^ microglia, decreased the proportion of DCX+ cells (8.1±0.6% vs 5.1±0.2% vs 9.0±0.7%, main effects of genotype F_(2,9)_=13.82, p=0.002, fig 4N) with post hoc analysis confirming wildtype microglia medium stimulated conditions were significantly different from control and *Cyfip1*^+/-^ stimulated conditions (p=0.008 and p=0.002, respectively). Similarly, only conditioned medium from wildtype microglia was associated with a significant increase in the proportion of apoptotic DCX+ cells (12.8±2.3% vs 28.3±5.0% vs 13.0±0.9%, main effects of genotype F_(2,9)_=6.12, p=0.021, fig 4O) with post hoc analysis confirming the wildtype stimulated condition was significantly different from the unconditioned medium controls and *Cyfip1*^+/-^ stimulated condition (p=0.007 and p=0.005, respectively). As before, no effect of any of the medium conditions was seen on the nestin-positive population (64.7±2.6% vs 64.0±2.3% vs 57.5±3.3%, F_(2,9)_=2.13, p=0.18, fig 4P). These data provide converging evidence for induction of neuronal apoptosis through factors secreted by microglia a process which is disrupted in *Cyfip1*^+/-^ animals.

### Proliferative cells are abnormally positioned in the *Cyfip1*^+/-^ hippocampus

As well as having to survive, newborn cells have to migrate to the correct location to functionally integrate into existing hippocampal circuitry. We noted that the gross positioning of new born cells appeared different in the *Cyfip1*^+/-^ animals. Cells undergoing proliferation are normally largely limited to the subgranular zone (SGZ) of the dentate gyrus, apart from some radially migrating neuroblasts undergoing their final divisions. Indeed, the majority of Ki67+ cells were found in the SGZ, however a significantly larger fraction was found in the SGZ in the *Cyfip1*^+/-^ animals (fig 5A,B, 80.4±1.1% vs 88.2±0.6%, t_(14)_=6.001, p= 3.2×10^-5^). As well as an increased fraction remaining in the SGZ, Cyfip1^+/-^ Ki67+ cells were also found significantly less far out into the granular zone (fig 5C, 17.0±1.4 μm vs 11.9±1.0 μm, t_(14)_=2.999, p= 0.009). These data suggested a migration deficit in *Cyfip1*^+/-^ animals.

**Figure 5:**
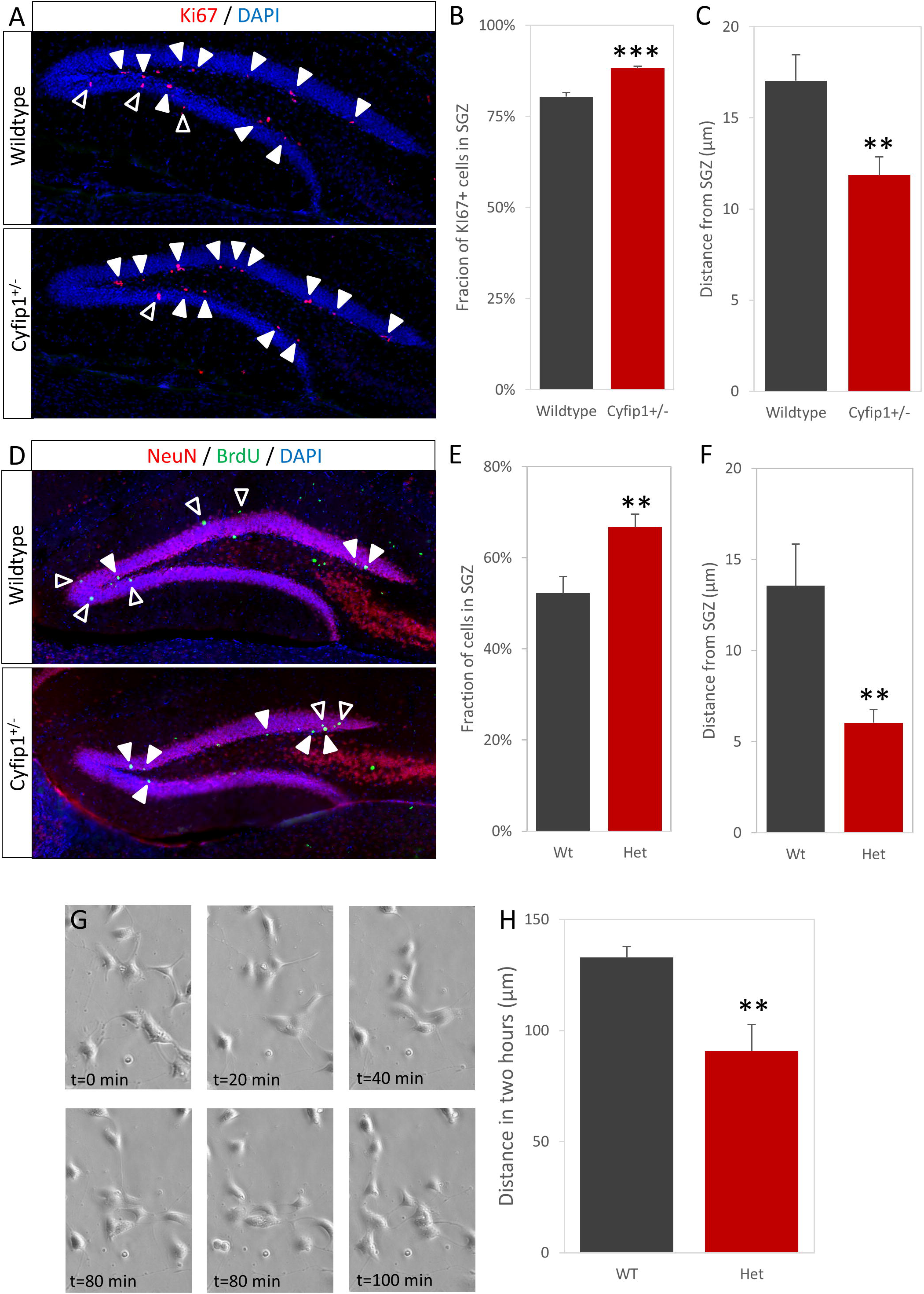
Altered migration of immature Cyfip1^+/-^ neurons. a) Immunohistochemistry for the Ki67, showing cells in the cell cycle, shows the majority are located in the SGZ (closed arrowheads), but some are starting to move out into the granular layer (open arrowheads). b) A significantly larger fraction of Ki67+ cells are located in the SGZ in *Cyfip1*^+/-^ animals, comparted to wild types. c) The distance moved from the SGZ is significantly lower in the *Cyfip1*^+/-^ animals, comparted to wild types. d) Post-mitotic adult born neurons (NeuN+/BrdU+) are located predominantly in the granular layer (open arrowheads), but some are still located in or near the SGZ (closed arrowheads). e) As with Ki67+ cells, a smaller fraction of *Cyfip1*^+/-^ cells are in the granular layer. f) Distance moved by these cells is significantly lower. g) Time-lapse imagining shows the movement of cells in primary hippocampal cultures. h) Quantification of cell movement during live imaging shows a significantly smaller distance moved by the *Cyfip1*^+/-^ cells.

### *Cyflpl*^+/-^ adult born neurons show impaired migration into the granular layer

In order to gain direct evidence of abnormal migration in *Cyfip1*^+/-^ animals we tracked the final positioning of newly born neurons using a 30 day BrdU pulse-chase paradigm followed by staining for BrdU/Neun. Neurons born in the SGZ normally undergo a radial migration outwards into the granular layer. Consistent with a migration deficit in the *Cyfip1*^+/-^ animals we found a greater proportion of BrdU+NeuN+ cells had remained in the SGZ (fig 5D,E, 52.2±3.6% vs 66.7±2.8%, t_(12)_=3.197, p= 0.008) and those cells that had migrated had moved a smaller distance into the granular zone (fig 5F, 13.6±2.3 μm vs 6.0±0.7 μm, t_(12)_=3.549, p= 0.004). We further analysed the location of the BrdU+/NeuN+ cells that had left the SGZ by dividing the dentate gyrus into four quartiles mapping on to the route taken by migrating cells, it was found that the distribution of cells over these quartiles was significantly different (fig S6, χ^2^=25.7, p=1.1×10^-5^) and, as would be anticipated with a migration deficit, BrdU+/NeuN+ cells in the *Cyfip1*^+/-^ animals were skewed towards the two most proximal quartiles (i.e. those closest to the starting point in the SGZ). Finally, we examined the putative migration phenotype observed *in vivo* by performing time-lapse imaging in primary hippocampal cultures. Over the two hour imaging period, *Cyfip1*^+/-^ cells covered significantly less distance than wildtype cells (fig 5G,H, 133.0±4.7 vs 90.7±12.0 μm, t_(10)_=3.993, p= 0.003). Both the *in vivo* and *in vitro* findings were consistent with a migration deficit due to *Cyfip1* haploinsufficiency.

### Actin dynamics are altered in *Cyfip1*^+/-^ cells and inhibition of Arp2/3 activity rescues migration deficits *in vitro*

As CYFIP1 regulates actin branching and polymerisation through the WAVE1 complex, a function crucial for cytoskeletal reorganisation and cell migration, we speculated that the *Cyyfp1*^+/-^ related migration phenotype was related to alterations in actin dynamics. A main determinant of actin function in the context of cytoskeleton reorganisation is the ratio of filamentous (F) to globular (G) actin. We first checked this ratio in primary hippocampal cultures, staining with fluorescently labelled phalloidin for F-actin and DNAse1 for G-actin, and found a significant increase in F-G actin ratio in *Cyfip1*^+/-^ cells (fig 6A,B, 0.67±0.3 vs 1.48±0.28, t_(5)_=2.904, p= 0.034). An increase in F-G actin ratio would be consistent with concurrent increased activity in Arp2/3 brought about by an expected reduction of CYFIP1-WAVE1 complexing under conditions of *Cyfip1* haploinsufficiency. Hence, we further speculated that treatment with Arp2/3 inhibitors would influence, potentially rescue, the abnormal cell migration phenotypes in Cyfip1^+/-^ cultures. To do this we used two drugs, CK-548, which induces a conformation change inhibiting actin binding and CK-666, which stabilises the inactive form of Arp2/3.

**Figure 6:**
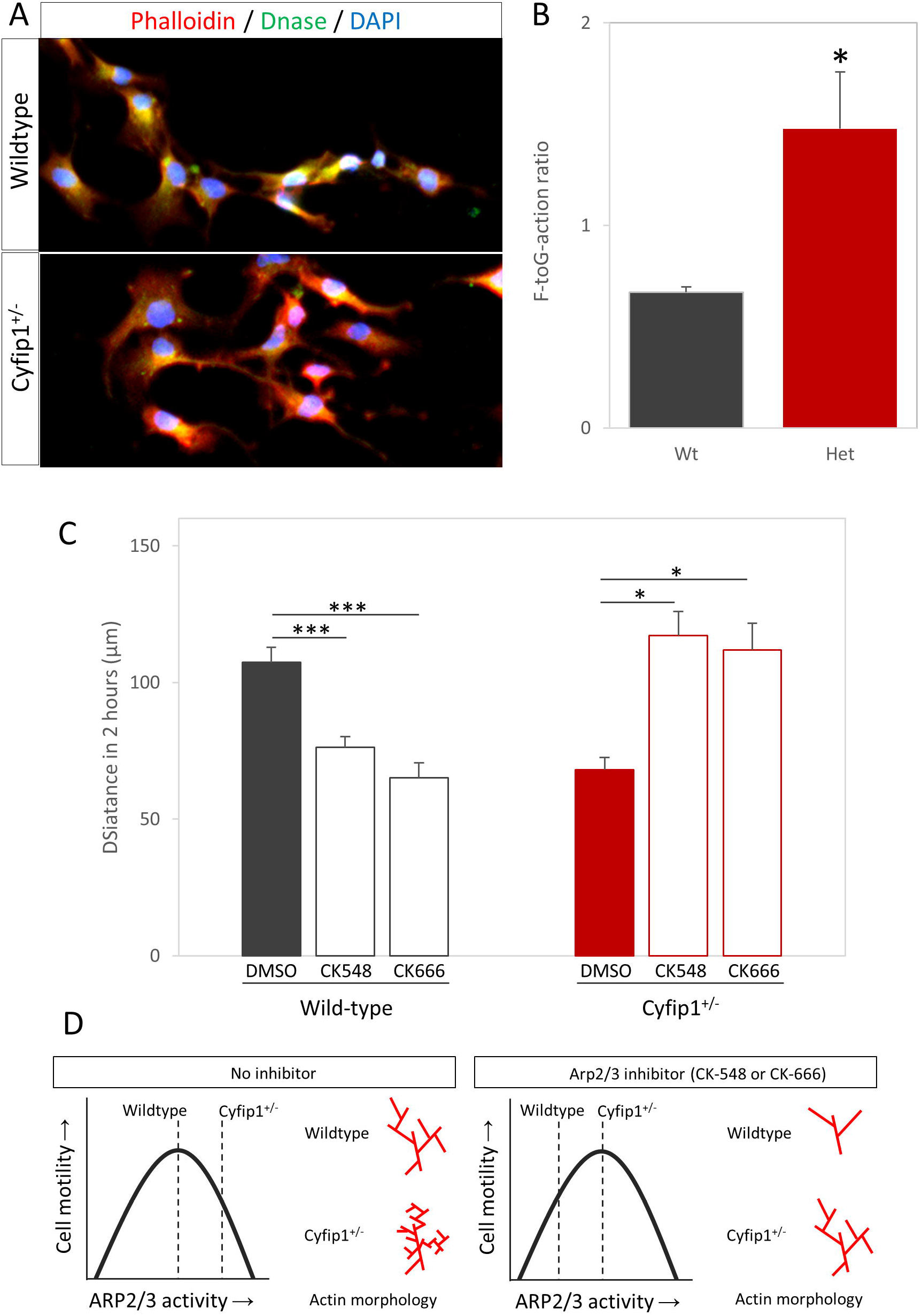
Disruption of Arp2/3 mediated actin dynamics is responsible for migration deficit. a) Fluorescently labelled phalloidin and DNAse show the amount of filamentous and globular actin, respectively. b) Quantificaion of fluorecene intensity shows *Cyfip1*^+/-^ have an increased F-to G-actin ratio, c) Treatment with Arp2/3 inhibitors CK-548 or CK-666 in wild-type cells significantly reduces the distance moved by cells during live imagine, whereas treatment of *Cyfip1*^+/-^ cells restoes distances to would-type levels. d) Model for effects of *Cyfip1* haploinsufficiency. At baseline conditions, *Cyfip1*^+/-^ cells have abnormally high Arp2/3 activity, leading to over-branching of the actin, and impaired cell motility. Following treatment with Arp2/3 inhibitors, the motility of wildtype cells is impaired, due to insufficient actin branching and polymerisation, but motility of Cyfip1^+/-^ is restored to normal levels.

Analysis by time-lapse showed that both drugs were effective in influencing cell migration. However, the effects were essentially opposite in wild type and Cyfip1^+/-^ cultures. In wild-type cultures, treatment with Arp2/3 inhibitors had a significant effect on migration (fig 6C, vehicle: 107.3±5.6, CK-548:76.3±3.9, CK-666: 65.2±5.4 μm, F_(2,24)_= 18.91, p=1.17×10^-5^), with both CK-548 and CK-666 showing significant decreases in distance moved (p=5.8×10^-4^ and 1.1×10^-5^, respectively). In *Cyfip1*^+/-^ cultures, there was also a significant effect on migration by Arp2/3 inhibitors (Vehicle: 68.1.3±4.4, CK-548:117.2±8.8, CK-666:112.0±9.8 μm, F_(2,6)_= 11.39, p=0.009). In contrast, both CK-548 and CK-666 caused significant increases in distance moved (p=0.011 and p=0.019, respectively), restoring migration to wild-type levels and suggesting excessive Arp2/3 activity was responsible for the migration deficit in *Cyfip1*^+/-^cells.

## Discussion

We have shown changes in the survival and migration of newly born neurons in hippocampus in a model of *Cyfip1* haploinsufficiency recapitulating the lowered gene dosage found in the pathogenic 15q11.2(BP1-BP2) copy number deletion. We report that *Cyfip1*^+/-^ animals have larger numbers of surviving newly born neurons in hippocampus, in the absence of increases in numbers of stem cells or rate of proliferation, and provide evidence that this is the result of a specific failure in microglia-induced apoptosis (see summary of proposed mechanism in fig S7). These newly born neurons also show an Arp2/3 dependent deficit in migration, resulting in a failure to migrate appropriately.

Our study is the first to our knowledge to demonstrate a direct effect of microglial soluble factors on the survival of immature neurons as a homeostatic mechanism. In contrast to earlier findings examining the effect of microglia on adult neurogenesis (Biscaro *et al*., 2012; Kohman *et al*., 2013; Mattei *et al*., 2014), we show this occurs in the absence of overt inflammatory disease or microglial activation. Hence, whilst *Cyfip1* is strongly up-regulated following activation with LPS, and it may play some role during the response to inflammatory stimuli, the phenotypes we observe are not dependent on any explicit pro-inflammatory stimulus. Our data indicate this is a mechanism susceptible to pathogenic changes however, as evidenced by the effects of *Cyfip1* haploinsufficiency on the ability of microglia to regulate apoptosis of immature neurons.

The effects of *Cyfip1* haploinsufficiency were highly cell-type specific and appeared limited to the stage of cell maturity where neuronal progenitors are fully committed to a neuronal phenotype, a stage of development characterised by high levels of apoptosis. Whilst we have demonstrated that soluble factors from microglia are involved in controlling numbers of immature neurons during adult neurogenesis, and that haploinsufficiency of *Cyfip1* leads to reduced apoptosis via an effect on factors secreted from microglia, the soluble factor(s) responsible remain to be identified. *Cyfip1* has pleotropic roles but the potential links between CYFIP1, actin remodelling, and cytokine secretion is especially interesting (Chemin *et al*., 2012). A focus for CYFIP1 action in microglia gains further evidence from a recent preprint demonstrating effects of cell specific deletion of Cyfip1 on microglia morphology, movement, and function (Drew *et al*., 2020).

In parallel, we have shown that adult born neurons in *Cyfip1*^+/-^ animals show a significantly impaired migration phenotype. Here our data point to a key role for CYFIP effects on actin dynamics mediated by Arp2/3 activity (see fig 6D). We hypothesise that lowered dosage will lead to reduced engagement of CYFIP protein with the CYFIP-WAVE1-Arp2/3 pathway resulting in reduced suppression of Arp2/3 activity and abnormal actin cytoskeleton remodelling required for migration. We evidence this by showing, as predicted, an increase in F-G actin ratio and rescue of the migration phenotype by pharmacological inhibition of Arp2/3 activity.

The combined effects of failure to undergo homeostatic apoptosis and failure to migrate to the correct location is likely to have profound effects on the functionality of adult born neurons in hippocampus, given the normal careful control of numbers and migration (Kempermann, Song and Gage, 2015). We observed lasting effects up to 30 days post division, when adult born neurons will already have local connections (Vivar *et al*., 2012). However, we cannot exclude the possibility that functional effects on brain and behaviour could be exerted at several points along the developmental pathway from the immature neuron stage to the fully integrated mature granule cell in the dentate gyrus. Haploinsufficiency of *Cyfip1* has been shown to have disease relevant behavioural consequences in both mice (Bachmann *et al*., 2019; Domínguez-Iturza *et al*., 2019) and rat (Silva *et al*., 2019) models, but to date, much of the neurobiology and behaviour in these models has remained unexplored. Our data would indicate a focus on hippocampal circuitry would prove of interest. Neuronal migration in the hippocampus has, to our knowledge, not been examined in patient populations. However, as AHN does take place in humans, and the biochemical pathways relevant to this study are well conserved, it is likely similar deficits would be observed in 15q11.2BP1-BP2 carriers.

The 15q11.2(BP1-BP2) deletion is not solely associated with schizophrenia, it also markedly increases risk for autism spectrum disorders (Chaste *et al*., 2014), ADHD and developmental delay (Abdelmoity *et al*., 2012), and epilepsy (de Kovel *et al*., 2010). Whether altered microglia functioning due to *Cyfip1* haploinsufficiency, impacting on adult hippocampal neurogenesis, also plays a part in these disorders is an open question, but all have been associated with alterations in immune system and microglia functioning (Jones and Thomsen, 2013; Tay *et al*., 2018). It is of note that a number of other psychiatric risk genes have also been shown to modify adult hippocampal neurogenesis in model systems. Whilst the effects are distinct from those of *Cyfip1*, each risk gene impacting on neurogenesis in its own way, this convergence of function may be indicative of a common pathogenic route.

## Acknowledgments

The authors would like to thank Chen Liang and Manal Adam for assistance with dissections, Caroline Best for assistance with genotyping, and Joe Davids for initial pilot immunohistochemistry. This work was supported by Wellcome Trust Strategic Award 100202/Z/12/Z (DEFINE) and the Hodge Foundation, UK. LJW is supported by a Waterloo Foundation Early Career Research Fellowship.

## Contributions

NH and LSW conceived and designed the study with input from WPG and JH. NH designed and performed experiments, JC performed animal husbandry/genotyping. LJW optimised and assisted with primary culture, WPG and LSW assisted with data interpretation, NH drafted the manuscript and MJO, WPG, JH, and LSW critically appraised and revised the manuscript. All authors approved the final manuscript

---

**Figure S1.**
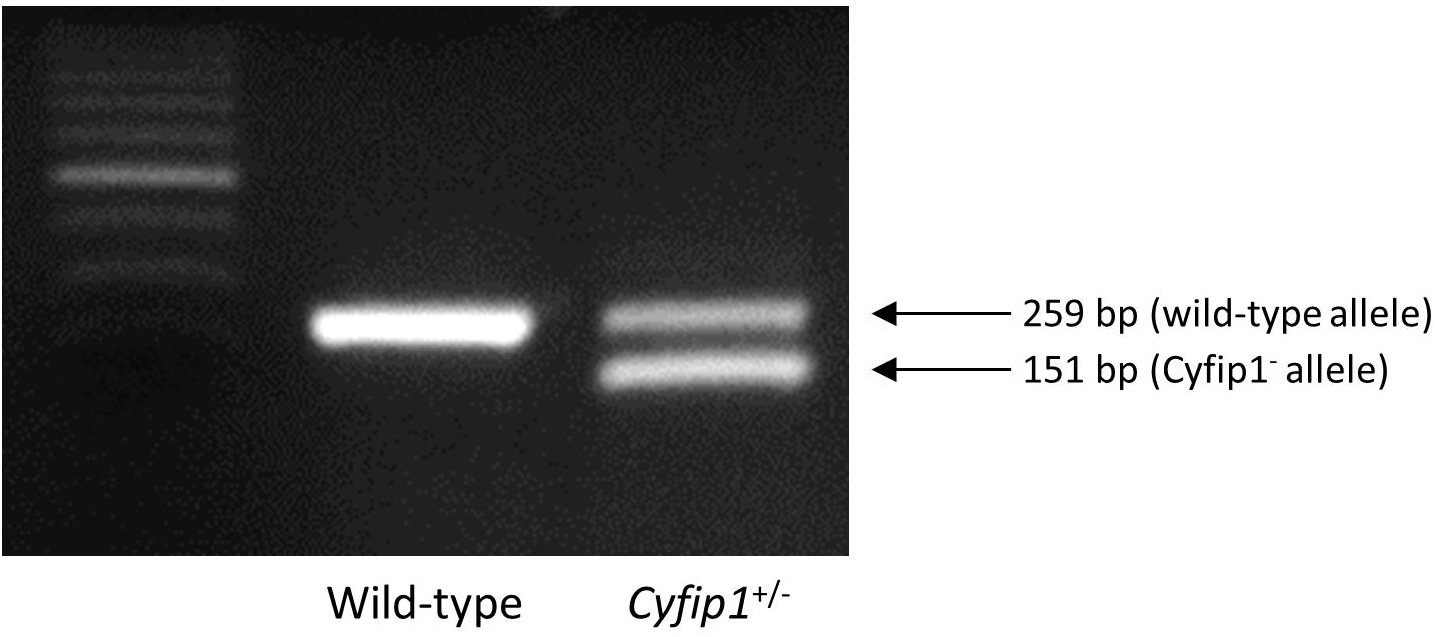

**Figure S2.**
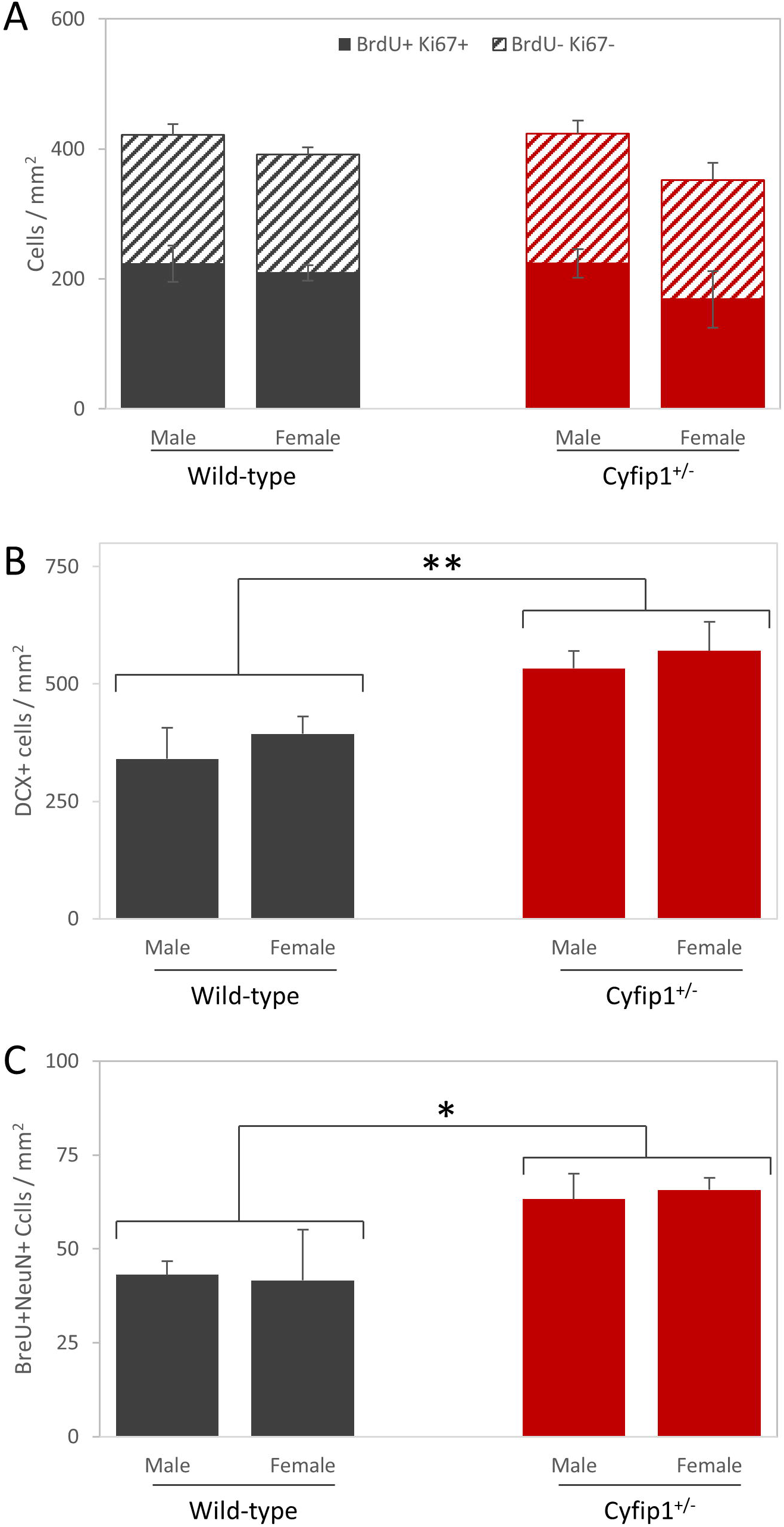

**Figure S3.**
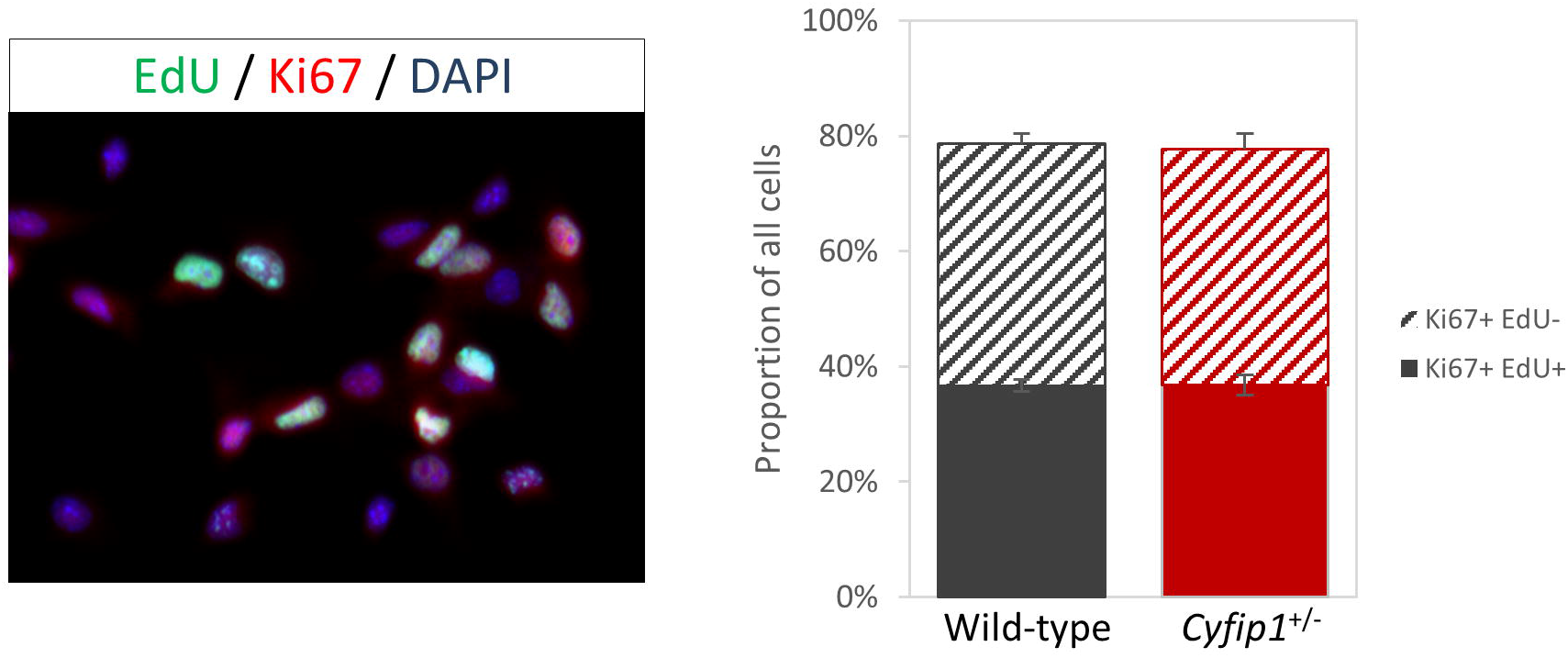

**Figure S4.**
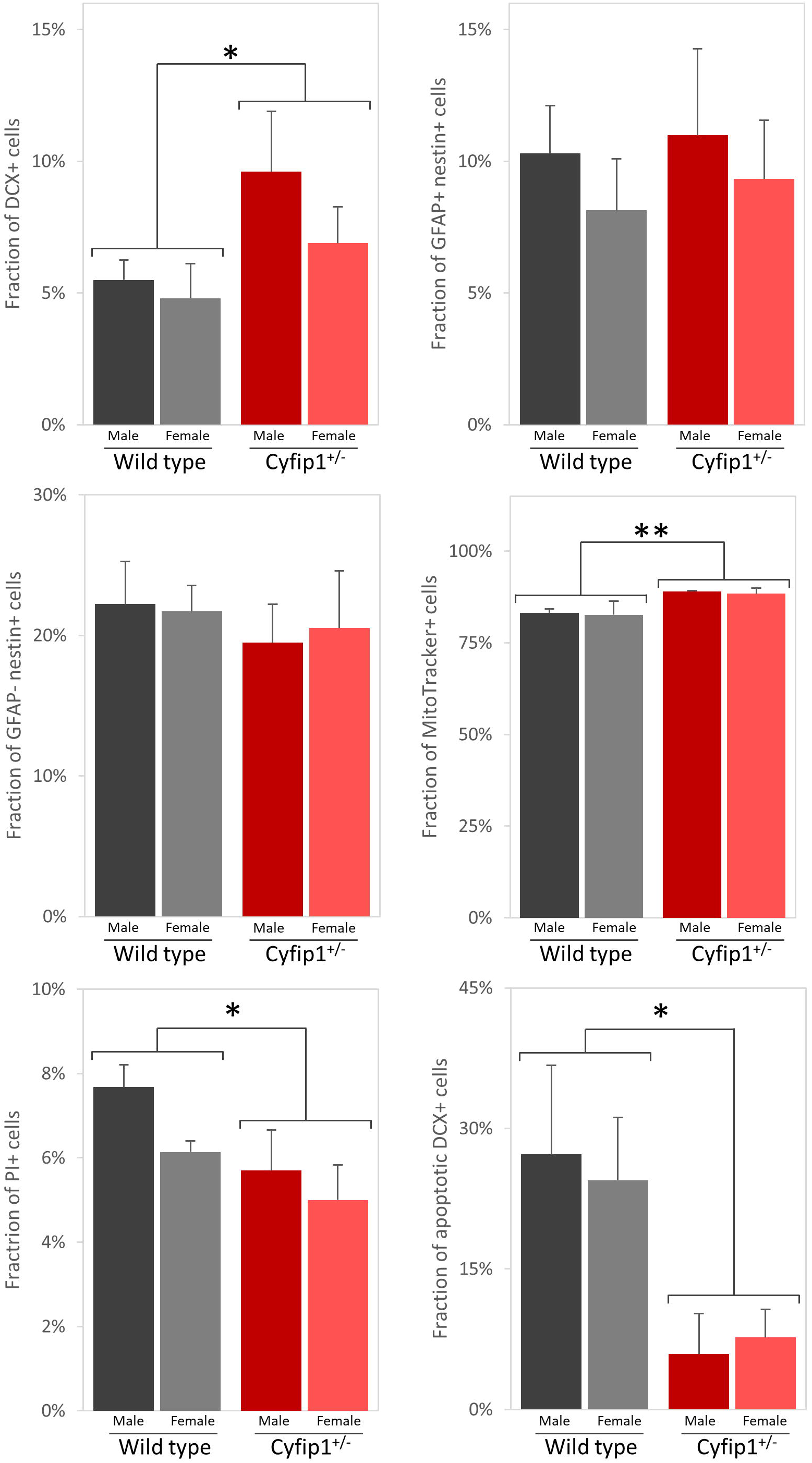

**Figure S5.**
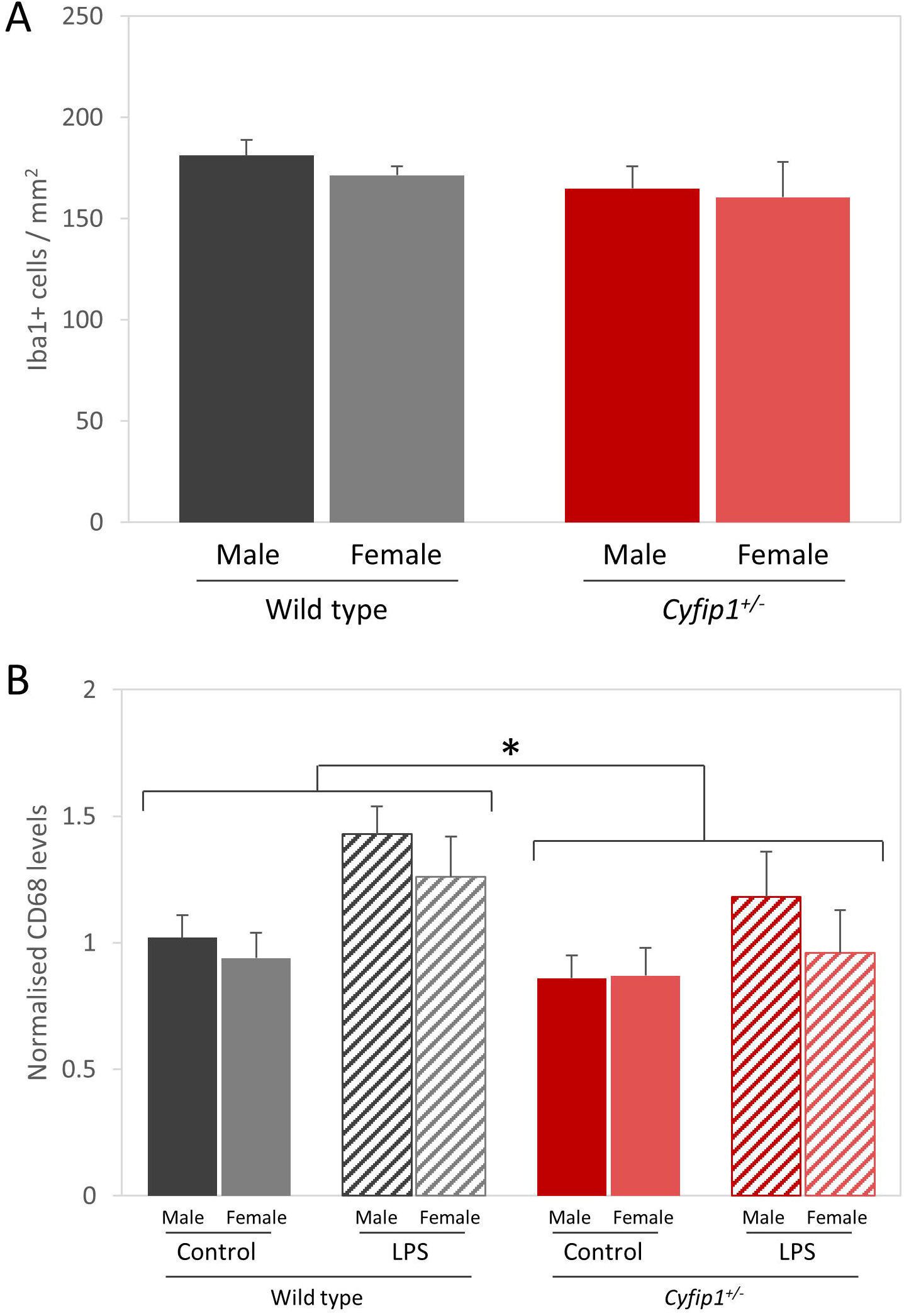

**Figure S6.**
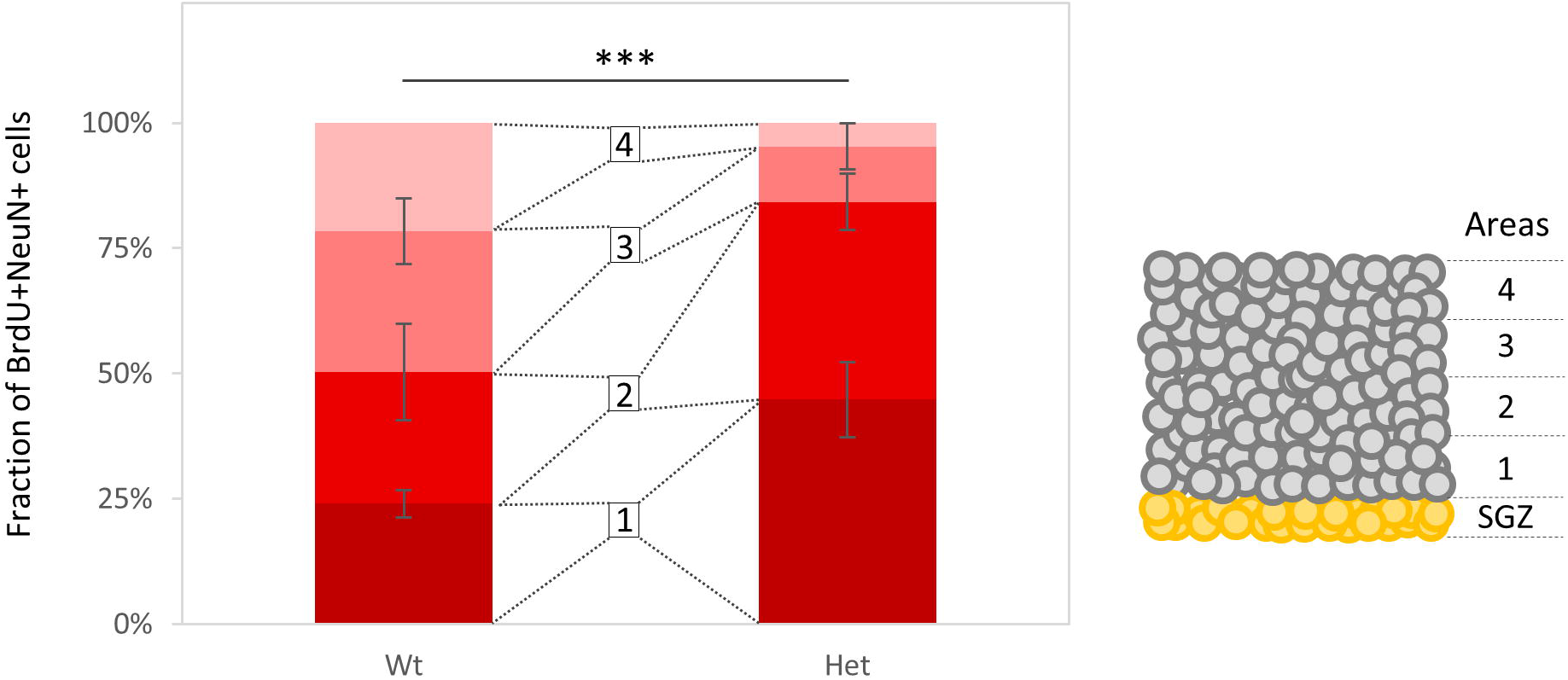

**Figure S7.**
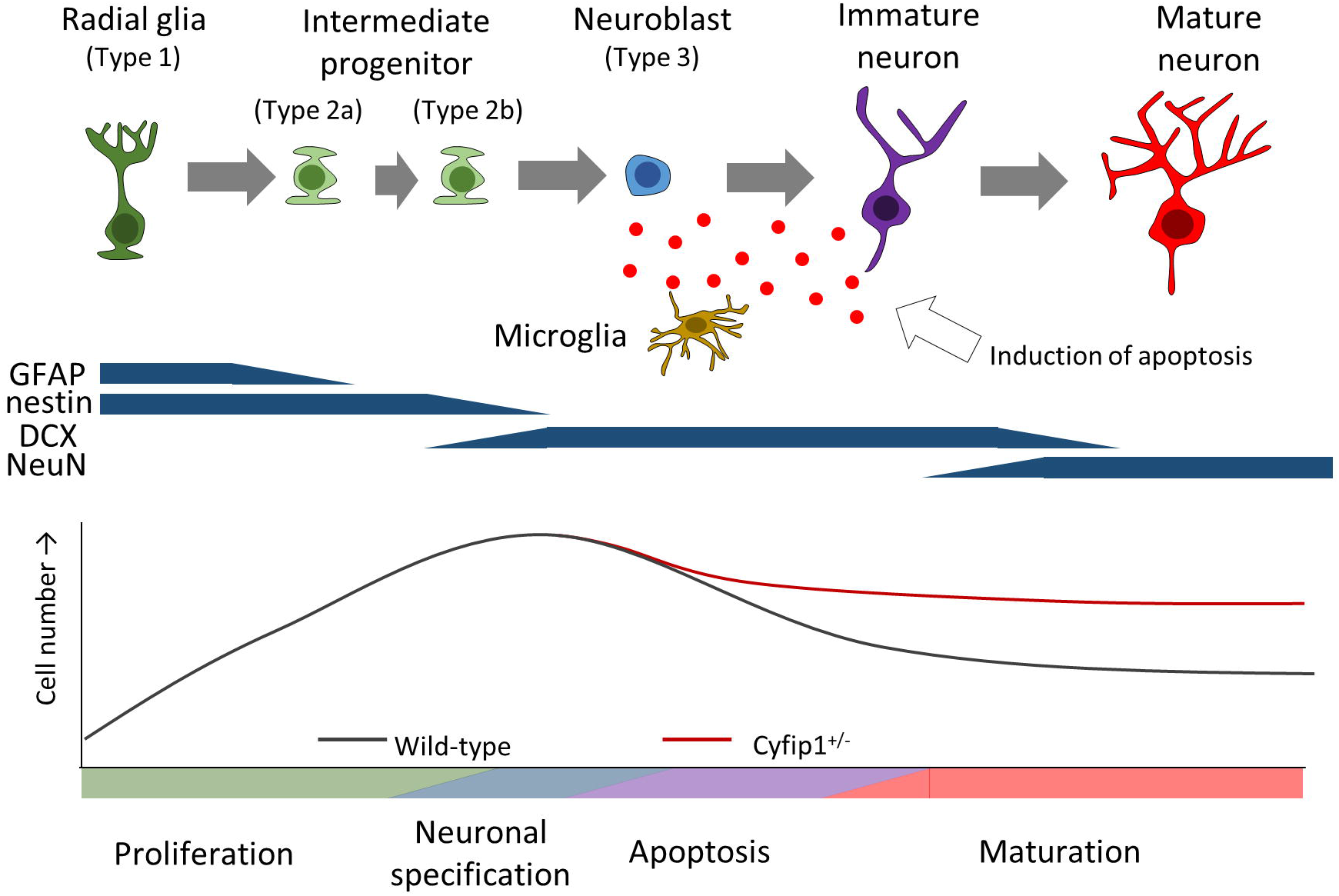

