## Supplementary information for "Haploinsufficiency of the psychiatric risk gene *Cyfip1* causes abnormal postnatal hippocampal neurogenesis through microglial and Arp2/3 mediated actin dependent mechanisms"

**Supplementary materials and methods**

*Cyfip1* heterozygous knockout mice

Cyfip1^tm2a(EUCOMM)Wtsi^ animals were obtained from the International Mouse Phenotyping Consortium (Harwell, UK) and maintained heterozygously on a C57BL6/J background. These mice have previously shown to have a significant reduction in brain *Cyfip1* expression(Pathania *et al.*, 2014). Groups used were of mixed sex. Animals were genotyped from ear or tail tissue. Genomic DNA was extracted using a Qiagen DNeasy Blood & Tissue kit, according to manufacturer’s protocol. Genotyping primers were 5’-TGGAAGTAATGGAACCGAACA, 5’-TCGTGGTATCGTTATGCGCC, and 5’-GTAACTACCTATAATGCAGACCTGAAG. See figure S1 for example of genotyping PCR products. All procedures were carried out according to the UK Animal (Scientific Procedures) Act 1986, Home Office Project Licence PPL 30/3135.

BrdU administration

*Cyfip1*^+/-^ animals and wildtype littermates were administered 100 mg/kg BrdU by IP injection from a 10 mg/ml stock solution in saline. 8 week old animals were either injected once and sacrificed 6 hour later or received five injections once daily, and sacrificed 30 days following first injection.

Primary hippocampal progenitor culture

Mixed hippocampal cultures were prepared from P7-P8 *Cyfip1*^+/-^ animals and wildtype littermates. Following cervical dislocation and removal of the brain, hippocampi were dissected and stored in ice cold PBS. Hippocampal tissue from individual animals was disrupted using a McIlwain tissue chopper and dissociated to single cells with 2 mg/ml papain (Sigma-Aldrich P4762) in standard medium (Neurobasal A medium (Thermo Fisher 10888022) containing 2% B27 supplement (Thermo Fisher 17504044), 1% Pen/Strep (Thermo Fisher 15240062) and 50 mM Glutamax (Thermo Fisher 35050061)), at 37 °C for 30 minutes. Following trituration in fresh standard medium, cell suspension was placed on top of layered 10 and 20% Optiprep (Sigma-Aldrich D1556) solutions in standard medium, and centrifuged at 600 rcf for 15 minutes.

The cell layer at the gradient interface was collected, washed in fresh standard medium, centrifuged at 250 rcf for 5 minutes, re-suspended in fresh standard medium and plated at 5x10^4^ cells on 13 mm glass coverslips, previously coated with 0.01% poly-l-lysine (Sigma-Aldrich P4707) and 10 µg/ml laminin (Sigma-Aldrich L2020). Cells were cultured at 37 °C, 5% CO_2_, 9% O_2_ with nitrogen for balance. Two hours following plating, all medium was replaced with fresh standard medium containing 20 ng/ml of both EGF and FGF2 (Sigma-Aldrich E9644 and SRP4037, respectively). Following three days of culture, 2/3rds of medium was replaced with fresh standard medium containing EGF and FGF2. Cultures were fixed at 5 DIV with ice cold 4% PFA for 30 minutes. Cell viability and death was assessed by incubation with 5 µg/ml DAPI (Sigma Aldrich D9542), 10 µg/ml propidium iodide (Sigma-Aldrich P3566) and 1 µM MitoTracker Green (Thermo Fisher M7514) for 40 minutes at 37 °C before cells were live-imaged.

Primary microglial culture

Primary microglia from whole brain were prepared from P7-P8 *Cyfip1*^+/-^ animals and wildtype littermates. Following cervical dislocation brains were removed and stored in ice cold PBS. Tissue from individual animals was disrupted using a McIlwain tissue chopper and dissociated to single cells with 0.25% Trypsin with 1 mM EDTA (Thermo Fisher 25200072) at 37 °C for 30 minutes. Dissociation was stopped by addition of glia medium (DMEM/F12 (Thermo Fisher 21331020) containing 10% heat inactivated fetal bovine serum (Thermo Fisher 10270106), 1% Pen/Strep and 200 mM Glutamax), and cells were centrifuged at 250 rcf for 5 minutes.

Following wash and centrifugation, cells were plated on previously poly-l-lysine coated plastic dishes, with one hemisphere plated on 20 cm^2^. Cells were cultured at 37 °C, 5% CO_2_, 9% O_2_ with nitrogen for balance. Medium was changed every third day until mixed glia were confluent at 7-10 days. Microglia were isolated by shake off at 250 rpm for 30 minutes, and cultured at 5x10^5^/ml on poly-l-lysine coated 13 mm glass coverslips or culture plastic as appropriate.

For production of conditioned medium, 24 hours following plating, microglia were changed to standard progenitor medium without growth factors and incubated for 24 hours. Medium was then collected and stored at -80 °C until use. Conditioned medium of different batches of cells was combined before use, to preclude batch-specific effects.

To assess activation, microglial cultures were stimulated with 10 μg/ml E.Coli lipopolysaccharide (Sigma-Aldrich L5293) for 24 hours prior to fixation or collection of medium.

Microglia depletion, microglia insert culture, and conditioned medium experiments

Microglia were depleted from primary cultures by addition of a single dose of 1 nM Mac-1-Sap (ATS Bio IT-06) with the medium change at 2 hours following isolation of cells. To study the effects of secreted factors from microglia on progenitor cultures, primary microglia were plated on hanging inserts (0.4 µm pore size Millicell inserts, Fisher Scientific PIHT12R48) following shake-off. Following 24 hours of culture in progenitor medium, inserts with 1.5x10^4^ microglia were added to progenitor cultures at 2 hours following progenitor isolation, and remained in the cultures until fixation. Control cultures had empty inserts. Microglia conditioned medium was added to progenitor medium at 25% at 2 hours following isolation, and was present for the duration of the experiment. Control conditions received 25% unconditioned medium.

Immunocytochemistry

Fixed cultures were blocked and permeabilized with 5% donkey serum and 0.1% Triton X100 in PBS for 30 minutes at room temperature. Primary antibodies were applied in the same solution overnight at 4 °C. Antibodies and detailed in Table S1. Following three washes in PBST, relevant secondary antibodies were applied in PBST for two hours at room temperature. For identification of microglia, Alexa 568 conjugated IB4 (Thermo Fisher I21412) was added to secondary antibody solutions at 2.5 µg/ml. Cells were washed, counterstained with DAPI, and mounted for microscopy.

| Table S1: antibodies used in the study | | | | |
| --- | --- | --- | --- | --- |
| **Target** | **Host** | **Manufacturer** | **Catalogue no** | **Dilution** |
| BrdU | Rt | Rio-Rad | OBT0030 | 1:500 |
| Ki-67 | Rb | Abcam | ab16667 | 1:1,000 |
| DCX | Gp | Millipore | AB2253 | 1:10,000 |
| NeuN | Ms | Millipore | MAB327 | 1:300 |
| Prox1 | Rb | Abcam | ab101851 | 1:500 |
| Nestin | Ms | Millipore | MAB353 | 1:500 |
| GFAP | Rt | Invitrogen | 13-0300 | 1:1,000 |
| Cleaved caspase 3 | Rb | CST | 9661S | 1:500 |
| Iba1 | Rb | Wako | 019-19741 | 1:2,000 |
| CD68 | Rt | Abcam | Ab5344 | 1:1,000 |

Image analysis

For sections, the entire dentate gyrus was acquired as a single image as a 20x magnification tile-scanned image. Stained cells were manually quantified by a genotype-blinded experimenter, and cell number is expressed as number of cells per mm^2^ of dentate gyrus.

For cultures, 8-12 randomly selected 20x fields of view were analysed per condition. Cells were manually quantified by a genotype-blinded experimenter, and cell number is expressed as fraction of total (DAPI^+^) cells unless otherwise noted. For microglial activation analysis, fluorescence intensity of CD68 and IB4 in individual cells was measured in ImageJ. CD68 fluorescence intensity was normalised to IB4 fluorescence intensity, and all values were then normalised to control conditions.

Quantitative PCR

RNA was isolated from primary microglia 24h following stimulation with LPS, and from unstimulated controls using the Qiagen RNeasy kit according to manufacturer’s protocol, DNAse treated and reverse transcribed with random hexamers. *Cyfip1* levels were determined by quantitative PCR, using 5’-TTCCTCTAGCATCTCGTTCA and 5’-ACCGTCCTCGCTGCTCTATC as primers. Data is expressed as fold change from control, normalised to GAPDH (5’-GAACATCATCCCTGCATCCA and 5’-CCAGTGAGCTTCCCGTTCA), according to the ΔΔCt method.

Statistical analysis and data availability

All n numbers represent individual animals, or cultures derived from individual animals. Group sized were determined based on previous experience. Appropriate statistical tests (two way Student T-test, one-way ANOVA with Tukey post-hoc test, or two-way ANOVA, as denoted in text) were employed in R, following tests for normality. Data is displayed as mean±SEM. P<0.05 is denoted by *, p<0.01 by ** and p<0.001 by ***. The data that support the findings of this study are available from the corresponding authors upon reasonable request.

**Supplementary results**

*In vitro* proliferation is unaffected by *Cyfip1* haploinsufficiency

In order to check if the lack of difference in proliferation seen in the hippocampus extends to our primary culture model system, we pulsed cultures with the nucleotide analogue EdU for 6 hours, and stained for Ki67 (fig S2A). There were no differences in the total number of Ki67+ cells in the cell cycle (78.7±1.7% vs 77.8±2.7%, t_(22)_=0.31 p=0.76) or the number of cells taking up EdU (36.7±1.0% vs 36.8±1.7%, t_(22)_=0.07 p=0.94) during the 6 hour pulse (fig S2B).

No sex differences in neurogenesis phenotype

As there have been reports of sexual dimorphism in neurogenesis phenotypes, we split our data by gender to investigate this. Looking at the levels of BrdU incorporation in the dentate gyrus after a 6 hour pulse (fig S3A), there are no significant effects of genotype (F_(1,7)_=0.68, p=0.45), sex (F_(1,7)_=0.003, p=0.96) or a significant interaction (F_(1,7)_=0, p=0.99). Similarly, there are no effects of genotype (F_(1,7)_=1.48, p=0.29), sex (F_(1,7)_=0.691, p=0.45) or a significant interaction (F_(1,7)_=0.76, p=0.43) on the number of Ki67+ cells.

When comparing the density of immature neurons in the dentate gyrus (fig S3B), there is a main effect of genotype (F_(1,13)_=13.05, p=0.005), but no main effect of sex (F_(1,13)_=0.79, p=0.40) or interaction (F_(1,13)_=0.025, p=0.89). Lastly, when looking at the numbers of BrdU+NeuN+ following a 30 d pulse-chase protocol separated by sex (fig S3C), there is a main effect of genotype (F_(1,12)_=8.98, p=0.01), but no main effect of sex (F_(1,12)_=0.004, p=0.95) or interaction (F_(1,12)_=0.07, p=0.80).

No sex differences in primary cultures

Numbers of immature DCX+ cells remain significantly increased in  *Cyfip1*^+/-^ animals when data is split by sex (fig S4A, with a significant effect of genotype (F_(1,12)_=7.59, p=0.012) and no significant effects of sex (F_(1,12)_=0.43, p=0.52) or a significant interaction (F_(1,12)_=0.64, p=0.44). The fraction of GFAP+ nestin+ Type 1/2a cells showed no sexual dimorphism (fig S4B), with no significant effects of sex (F_(1,32)_=0.79, p=0.38), genotype (F_(1,32)_=0.15, p=0.71) or a significant interaction (F_(1,34)_=0.02, p=0.88). Likewise, there were no sex differences in the fraction of GFAP- nestin+ cells Type 2b (fig S4C) , with no significant effects of sex (F_(1,32)_=0, p=0.98), genotype (F_(1,32)_=0.43, p=0.52) or a significant interaction (F_(1,34)_=0.06, p=0.81).

As seen previously, there were no sex differences in cell viability (fig S4D), showing a significant effect of genotype (F_(1,10)_=13.33, p=0.004), but not of sex F_(1,10)_=0.55, p=0.47, or a significant interaction (F_(1,10)_=0.14, p=0.71). Similarly, the number of dead/dying cells marked by PI (fig S4E) is significantly affected by genotype (F_(1,11)_=8.59, p=0.014), but not sex (F_(1,11)_=2.18, p=0.17), without any significant interaction (F_(1,11)_=0.30, p=0.60). Lastly, the fraction of DCX+ cells which are apoptotic (fig S4F) is also unaffected by sex (F_(1,13)_=0.32, p=0.58), but is significantly affected genotype (F_(1,13)_=4.91, p=0.04), in the absence of an interaction effect (F_(1,13)_=0.03, p=0.87).

Effects of *Cyfip1* on microglia are not sex dependent

When investigating the density of microglia in the hippocampus of *Cyfip1^+/-^* animals and wildtype littermates (fig S7A), we found no effect of sex, with no main effects of sex (F_(1,12)_=0.35 p=0.59), genotype (F_(1,12)_=1.05 p=0.332), or a significant interaction (F_(1,12)_=0.05 p=0.83). When assessing the effects of genotype and sex on microglial activation (fig S7B), we found significant effects of genotype (F_(1,59)_=4.82, p=0.033) and LPS stimulation (F_(1,59)_=8.179, p=0.006, but not of sex (F_(1,59)_=1.300, p=0.26).

Figure S1 *– Representative genotyping image for the Cyfip1^+/-^ mouse line.*

Figure S2: *In vivo neurogenesis phenotype split by sex*. a) Analysis of proliferation indexed by a 6 h BrdU pulse with Ki67 immunohistochemistry show no effects of sex or genotype. b) Immunohistochemistry for the immature neuronal marker DCX show a significant effect of genotype in the absence of any effects of sex. c) Analysis of mature adult born neurons, by NeuN immunohistochemistry 30 days after BrdU pulsing, showed a significant effect of genotype, again in the absence of any sex effects.

Figure S3: *Proliferation in primary hippocampal cultures*. a) Following withdrawal of growth factors, cells were given a 6 h EdU pulse and stained for Ki67. b) As seen following the *in vivo* BrdU pulse experiments, there are no differences in the total number of Ki67+ cells in the cell cycle or the fraction that had undergone cell division (EdU+).

Figure S4: *Primary cultures split by sex.* a) The significant increase in DCX proportion seen in *Cyfip1*^+/-^ animals is not sexually dimorphic. b) There is no significant difference in the proportion of early nestin+ GFAP+ Type 1/2a cells, either by genotype or by sex. c) The fraction of nestin+ GFAP- Type 2b cells is unaffected by genotype or sex. d) The significant increase in the proportion of MitoTracker+ viable cells is not affected by sex. e) The sex of the animals has no effect on the significant decrease of the proportion of dead or dying PI+ cells. f) The fraction of DCX+ cells which are positive for the apoptotic marker cleaved caspase 3 is only affected by genotype, not by sex.

Figure S5: *Effects of sex on microglia.* a) Density of microglia in the dentate gyrus of the hippocampus is unaffected by either sex or genotype. b) Effects of genotype and LPS stimulation in primary microglia are not dependent on sex.

Figure S6: *Distribution of adult born neurons is different in Cyfip1^+/-^ animals.* When dividing the thickness of the dentate gyrus into four equal quartiles, NeuN+/BrdU+ cells in *Cyfip*^+/-^ animals were significantly more likely to be found in the two quartiles proximal to the SGZ, compared to the more even distribution in wildtype littermates.

Figure S7: *Model for effects of Cyfip1 haploinsufficiency*. Progression of cells through the different stages of neurogenesis is unaltered though the early radial glial and intermediate progenitor stages, with no changes in number of proliferation. However, when the doublecortin positive stage of development is reached, there is a failure of *Cyfip1*^+/-^ microglia to induce neuronal apoptosis, as takes place in wild-type animals. This leads to the survival of a significantly larger number of immature neurons into maturity.
